## Supporting Information for "An inductive bias for slowly changing features in human reinforcement learning"

‡SHM and NWS are joint senior authors.
* noa.hedrich [at] uni-hamburg.de
* nicolas.schuck [at] uni-hamburg.de

### Text A Preregistration

We submitted two preregistrations (see https://osf.io/6dy8f). The first was submitted before any data collection, the second before collecting the replication sample. The hypothesis, core tenets of the task (learning phase and test phase, bi-dimensional stimuli with slow/fast and relevant/irrelevant feature) and main analyses (t-tests) in the first preregistration are maintained. Yet, piloting led us to make changes to the task design and analysis to better address our research question and replicate our initial findings. Therefore, we deviated from the first preregistration. To validate our results from experiment 2, we updated the preregistration to reflect the changes to the task and analysis and then conducted a direct replication of experiment 2 in strict adherence to this preregistration. For consistency, all behavioural analyses reported in the main text are those specified in the second preregistration. The modelling analyses are not included in the preregistration.

The original preregistration lists mean accuracy below 65% as an exclusion criterion, however during data collection it became clear that this criterion was too stringent, as it would entail excluding 33 participants in experiment 1 and 20 participants in experiment 2. Therefore, this criterion was not applied, instead, no participants were excluded in these experiments. In the updated preregistration we specified a new exclusion criterion, which was applied to the replication sample (see Methods).

a)

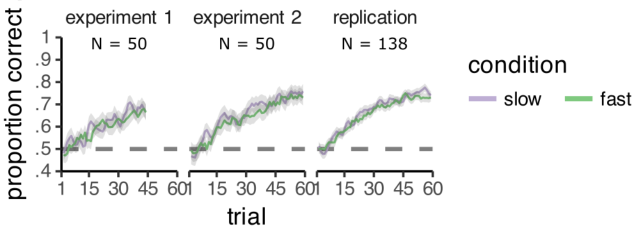


b)

| | | Slow Blocks | | Fast Blocks | | | |
|---|---|---|---|---|---|---|---|
| **Sample** | **N** | **M** | **SD** | **M** | **SD** | **Hedge's g (95% CI)** | **Weight** |
| Experiment 1 | 50 | 128.1 | 99.2 | 108.9 | 105.9 | 0.19 [-0.21; 0.58] | 21.0% |
| Experiment 2 | 50 | 248.6 | 152.3 | 217.6 | 158.6 | 0.20 [-0.19; 0.59] | 21.0% |
| Replication | 138 | 270.2 | 126.0 | 242.9 | 128.7 | 0.21 [-0.02; 0.45] | 58.0% |
| **Overall** | **238** | | | | | **0.20 [ 0.02; 0.38]** | **100.0%** |

Heterogeneity: $I^2 = 0\%$, $\tau^2 = 0$, $p = 0.99$

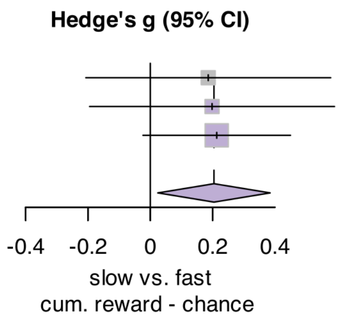


c)

| | | Slow Blocks | | Fast Blocks | | | |
|---|---|---|---|---|---|---|---|
| **Sample** | **N** | **M** | **SD** | **M** | **SD** | **Hedge's g (95% CI)** | **Weight** |
| Experiment 1 | 50 | 0.71 | 0.148 | 0.71 | 0.128 | -0.01 [-0.40; 0.38] | 21.0% |
| Experiment 2 | 50 | 0.76 | 0.114 | 0.74 | 0.106 | 0.20 [-0.20; 0.59] | 20.9% |
| Replication | 138 | 0.75 | 0.101 | 0.74 | 0.098 | 0.07 [-0.17; 0.31] | 58.0% |
| **Overall** | **238** | | | | | **0.08 [-0.10; 0.26]** | **100.0%** |

Heterogeneity: $I^2 = 0\%$, $\tau^2 = 0$, $p = 0.75$

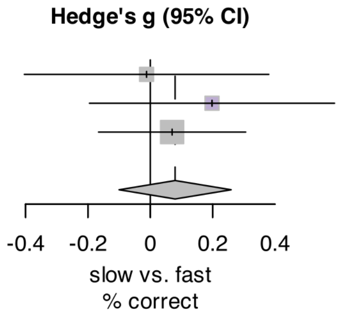


Figure A: **Participant learning curves and meta analysis. A:** Participant learning curves for the slow and fast condition, showing the increase in the accuracy of choices across trials of the learning phase in the slow (purple) and fast(green) blocks. Each panel shows a different experiment sample. **B:** Results of the meta analysis of the cumulative reward in the learning phase, confirming consistently higher reward in the slow blocks. **C:** Results of the meta analysis of the accuracy in the test phase, showing no consistent advantage in generalisation in slow blocks. Plots produced with the `forest` function from the `meta` package in R [1].

### Effect sizes and confidence intervals for the mixed effects models

The following tables report the parameter estimates for the mixed effect models. $\mathrm{CI}_l$ and $\mathrm{CI}_u$ denote the lower and upper bound of the 95% confidence interval, respectively. The standard deviation of the random effects is reported under $\sigma_{\mathrm{rand}}$. Not all fixed effects are included as random effects, if there were issues fitting the full random effects. For all models, random effects are per subject. If model comparison revealed the need to remove the effect of interest, both the model with the effect of interest reported in the main text and the best model with all non-significant terms removed are reported.

| Parameter | Estimate | $\mathrm{CI}_l$ | $\mathrm{CI}_u$ | $\sigma_{\mathrm{rand}}$ |
|---|---|---|---|---|
| (Intercept) | 3.04 | 2.49 | 3.59 | 1.77 |
| condition [slow] | 0.50 | 0.17 | 0.83 | 0.30 |
| $R_t$ | -0.19 | -0.40 | 0.02 | 0.45 |
| trial | -2.42 | -3.08 | -1.76 | 2.08 |
| condition$\times R_t$ | 0.09 | 0.02 | 0.17 | |
| condition$\times$ trial | -0.76 | -1.17 | -0.34 | |
| trial$\times R_t$ | 1.13 | 0.92 | 1.35 | |

Table A: **Experiment 1: Best model predicting participant learning phase choices.**

| Parameter | Estimate | $\mathrm{CI}_l$ | $\mathrm{CI}_u$ | $\sigma_{\mathrm{rand}}$ |
|---|---|---|---|---|
| (Intercept) | 3.20 | 2.74 | 3.67 | 1.47 |
| condition [slow] | 0.24 | -0.08 | 0.55 | 0.38 |
| $R_t$ | -0.56 | -0.79 | -0.34 | 0.57 |
| trial | -2.59 | -3.16 | -2.01 | 1.83 |
| condition [slow]$\times R_t$ | 0.12 | 0.05 | 0.19 | |
| condition [slow]$\times$ trial | -0.36 | -0.72 | -0.00 | |
| trial$\times R_t$ | 1.81 | 1.62 | 2.00 | |

Table B: **Experiment 2: Best model predicting participant learning phase choices.**

| Parameter | Estimate | $\mathrm{CI}_l$ | $\mathrm{CI}_u$ | $\sigma_{\mathrm{rand}}$ |
|---|---|---|---|---|
| (Intercept) | 3.59 | 3.29 | 3.89 | 1.59 |
| condition [slow] | 0.24 | 0.05 | 0.43 | 0.32 |
| $R_t$ | -0.48 | -0.61 | -0.35 | 0.48 |
| trial | -3.14 | -3.46 | -2.82 | 1.66 |
| condition$\times R_t$ | 0.11 | 0.06 | 0.15 | |
| condition$\times$ trial | -0.35 | -0.57 | -0.13 | |
| trial$\times R_t$ | 1.80 | 1.68 | 1.92 | |

Table C: **Replication: Best model predicting participant learning phase choices.**

| Parameter | Estimate | $CI_l$ | $CI_u$ | $\sigma_{\text{rand}}$ |
|---|---|---|---|---|
| (Intercept) | 2.94 | 2.64 | 3.25 | 0.74 |
| $cos(\theta_R)$ | -0.65 | -0.96 | -0.34 | |
| $sin(\theta_R)$ | 0.07 | -0.25 | 0.38 | |
| $cos(\theta_I)$ | 0.26 | -0.05 | 0.58 | |
| $sin(\theta_I)$ | -0.00 | -0.32 | 0.31 | |
| trial | -2.49 | -2.78 | -2.19 | |
| $cos(\theta_R)$ ×trial | 2.16 | 1.75 | 2.58 | |
| $sin(\theta_R)$ ×trial | -0.03 | -0.44 | 0.39 | |
| $cos(\theta_I)$ ×trial | -0.41 | -0.82 | 0.00 | |
| $sin(\theta_I)$ ×trial | 0.03 | -0.39 | 0.44 | |

Table D: **Experiment 1: Mixed effects model predicting participant learning phase choices from the feature positions in the slow blocks.**

| Parameter | Estimate | $CI_l$ | $CI_u$ | $\sigma_{\text{rand}}$ |
|---|---|---|---|---|
| (Intercept) | 2.53 | 2.25 | 2.81 | 0.68 |
| $cos(\theta_R)$ | -0.67 | -0.97 | -0.38 | |
| $sin(\theta_R)$ | -0.17 | -0.46 | 0.12 | |
| $cos(\theta_I)$ | -0.36 | -0.65 | -0.06 | |
| $sin(\theta_I)$ | -0.08 | -0.38 | 0.21 | |
| trial | -1.83 | -2.10 | -1.55 | |
| $cos(\theta_R)$ ×trial | 2.06 | 1.67 | 2.45 | |
| $sin(\theta_R)$ ×trial | 0.20 | -0.19 | 0.58 | |
| $cos(\theta_I)$ ×trial | 0.46 | 0.07 | 0.85 | |
| $sin(\theta_I)$ ×trial | 0.13 | -0.26 | 0.52 | |

Table E: **Experiment 1: Mixed effects model predicting participant learning phase choices from the feature positions in the fast blocks.**

| Parameter | Estimate | $CI_l$ | $CI_u$ | $\sigma_{\text{rand}}$ |
|---|---|---|---|---|
| (Intercept) | 3.02 | 2.75 | 3.29 | 0.62 |
| $cos(\theta_R)$ | -0.89 | -1.18 | -0.60 | |
| $sin(\theta_R)$ | -0.06 | -0.35 | 0.22 | |
| $cos(\theta_I)$ | -0.02 | -0.31 | 0.27 | |
| $sin(\theta_I)$ | 0.04 | -0.24 | 0.33 | |
| trial | -2.49 | -2.74 | -2.24 | |
| $cos(\theta_R)$ ×trial | 2.73 | 2.37 | 3.09 | |
| $sin(\theta_R)$ ×trial | 0.06 | -0.28 | 0.41 | |
| $cos(\theta_I)$ ×trial | -0.05 | -0.40 | 0.30 | |
| $sin(\theta_I)$ ×trial | -0.03 | -0.38 | 0.32 | |

Table F: **Experiment 2: Mixed effects model predicting participant learning phase choices from the feature positions in the slow blocks.**

| Parameter | Estimate | $\text{CI}_l$ | $\text{CI}_u$ | $\sigma_{\text{rand}}$ |
|---|---|---|---|---|
| (Intercept) | 2.87 | 2.60 | 3.14 | 0.66 |
| $cos(\theta_R)$ | -1.16 | -1.43 | -0.88 | |
| $sin(\theta_R)$ | -0.18 | -0.46 | 0.10 | |
| $cos(\theta_I)$ | -0.17 | -0.44 | 0.11 | |
| $sin(\theta_I)$ | -0.09 | -0.37 | 0.19 | |
| trial | -2.22 | -2.47 | -1.98 | |
| $cos(\theta_R)$ ×trial | 2.90 | 2.56 | 3.25 | |
| $sin(\theta_R)$ ×trial | 0.20 | -0.14 | 0.54 | |
| $cos(\theta_I)$ ×trial | 0.08 | -0.26 | 0.42 | |
| $sin(\theta_I)$ ×trial | 0.11 | -0.23 | 0.45 | |

Table G: **Experiment 2: Mixed effects model predicting participant learning phase choices from the feature positions in the fast blocks.**

| Parameter | Estimate | $\text{CI}_l$ | $\text{CI}_u$ | $\sigma_{\text{rand}}$ |
|---|---|---|---|---|
| (Intercept) | 4.07 | 3.71 | 4.44 | 1.93 |
| $cos(\theta_R)$ | -0.47 | -0.72 | -0.23 | 0.84 |
| $sin(\theta_R)$ | 0.03 | -0.17 | 0.22 | 0.19 |
| $cos(\theta_I)$ | -0.08 | -0.27 | 0.11 | 0.23 |
| $sin(\theta_I)$ | 0.09 | -0.10 | 0.29 | 0.22 |
| trial | -3.73 | -4.11 | -3.34 | 2.01 |
| $cos(\theta_R)$ ×trial | 2.57 | 2.33 | 2.82 | |
| $sin(\theta_R)$ ×trial | -0.01 | -0.23 | 0.22 | |
| $cos(\theta_I)$ ×trial | 0.01 | -0.22 | 0.23 | |
| $sin(\theta_I)$ ×trial | -0.11 | -0.33 | 0.12 | |

Table H: **Replication: Mixed effects model predicting participant learning phase choices from the feature positions in the slow blocks.**

| Parameter | Estimate | $\text{CI}_l$ | $\text{CI}_u$ | $\sigma_{\text{rand}}$ |
|---|---|---|---|---|
| (Intercept) | 3.57 | 3.29 | 3.85 | 1.44 |
| $cos(\theta_R)$ | -0.71 | -0.93 | -0.48 | 0.71 |
| $sin(\theta_R)$ | -0.05 | -0.24 | 0.14 | 0.26 |
| $cos(\theta_I)$ | -0.10 | -0.29 | 0.09 | 0.18 |
| $sin(\theta_I)$ | -0.04 | -0.23 | 0.14 | 0.11 |
| trial | -3.09 | -3.40 | -2.78 | 1.54 |
| $cos(\theta_R)$ ×trial | 2.62 | 2.39 | 2.85 | |
| $sin(\theta_R)$ ×trial | 0.07 | -0.15 | 0.29 | |
| $cos(\theta_I)$ ×trial | 0.08 | -0.14 | 0.30 | |
| $sin(\theta_I)$ ×trial | 0.04 | -0.18 | 0.26 | |

Table I: **Replication: Mixed effects model predicting participant learning phase choices from the feature positions in the fast blocks.**

| Parameter | Estimate | $\text{CI}_l$ | $\text{CI}_u$ | $\sigma_{\text{rand}}$ |
|---|---|---|---|---|
| (Intercept) | 0.01 | -0.08 | 0.10 | 0.15 |
| condition [slow] | 0.06 | -0.05 | 0.17 | |
| $R_{\text{diff},t}$ | 1.01 | 0.92 | 1.10 | |
| condition [slow] × $R_{\text{diff},t}$ | -0.02 | -0.14 | 0.11 | |

Table J: **Experiment 1: Mixed effects model predicting participant test phase choices.**

| Parameter | Estimate | $\text{CI}_l$ | $\text{CI}_u$ | $\sigma_{\text{rand}}$ |
|---|---|---|---|---|
| (Intercept) | 0.04 | -0.03 | 0.11 | 0.15 |
| $R_{\text{diff},t}$ | 1.00 | 0.94 | 1.06 | |

Table K: **Experiment 1: Best model predicting participant test phase choices.**

| Parameter | Estimate | $\mathrm{CI}_l$ | $\mathrm{CI}_u$ | $\sigma_{\mathrm{rand}}$ |
|---|---|---|---|---|
| (Intercept) | 0.06 | -0.02 | 0.13 | 0.20 |
| condition [slow] | -0.04 | -0.12 | 0.04 | 0.10 |
| $R_{\mathrm{diff},t}$ | 1.19 | 1.13 | 1.25 | |
| condition [slow] $\times R_{\mathrm{diff},t}$ | 0.13 | 0.04 | 0.21 | |

Table L: **Experiment 2: Best model predicting participant test phase choices.**

| Parameter | Estimate | $\mathrm{CI}_l$ | $\mathrm{CI}_u$ | $\sigma_{\mathrm{rand}}$ |
|---|---|---|---|---|
| (Intercept) | -0.03 | -0.06 | 0.01 | 0.12 |
| condition [slow] | 0.01 | -0.04 | 0.06 | 0.06 |
| $R_{\mathrm{diff},t}$ | 1.36 | 1.23 | 1.49 | 0.73 |
| condition [slow] $\times R_{\mathrm{diff},t}$ | 0.09 | -0.02 | 0.21 | 0.57 |

Table M: **Replication: Mixed effects model predicting participant test phase choices.**

| Parameter | Estimate | $\mathrm{CI}_l$ | $\mathrm{CI}_u$ | $\sigma_{\mathrm{rand}}$ |
|---|---|---|---|---|
| (Intercept) | -0.02 | -0.05 | 0.01 | 0.12 |
| condition [slow] | | | | 0.06 |
| $R_{\mathrm{diff},t}$ | 1.40 | 1.28 | 1.52 | 0.74 |
| condition [slow] $\times R_{\mathrm{diff},t}$ | | | | 0.57 |

Table N: **Replication: Best model predicting participant test phase choices.**

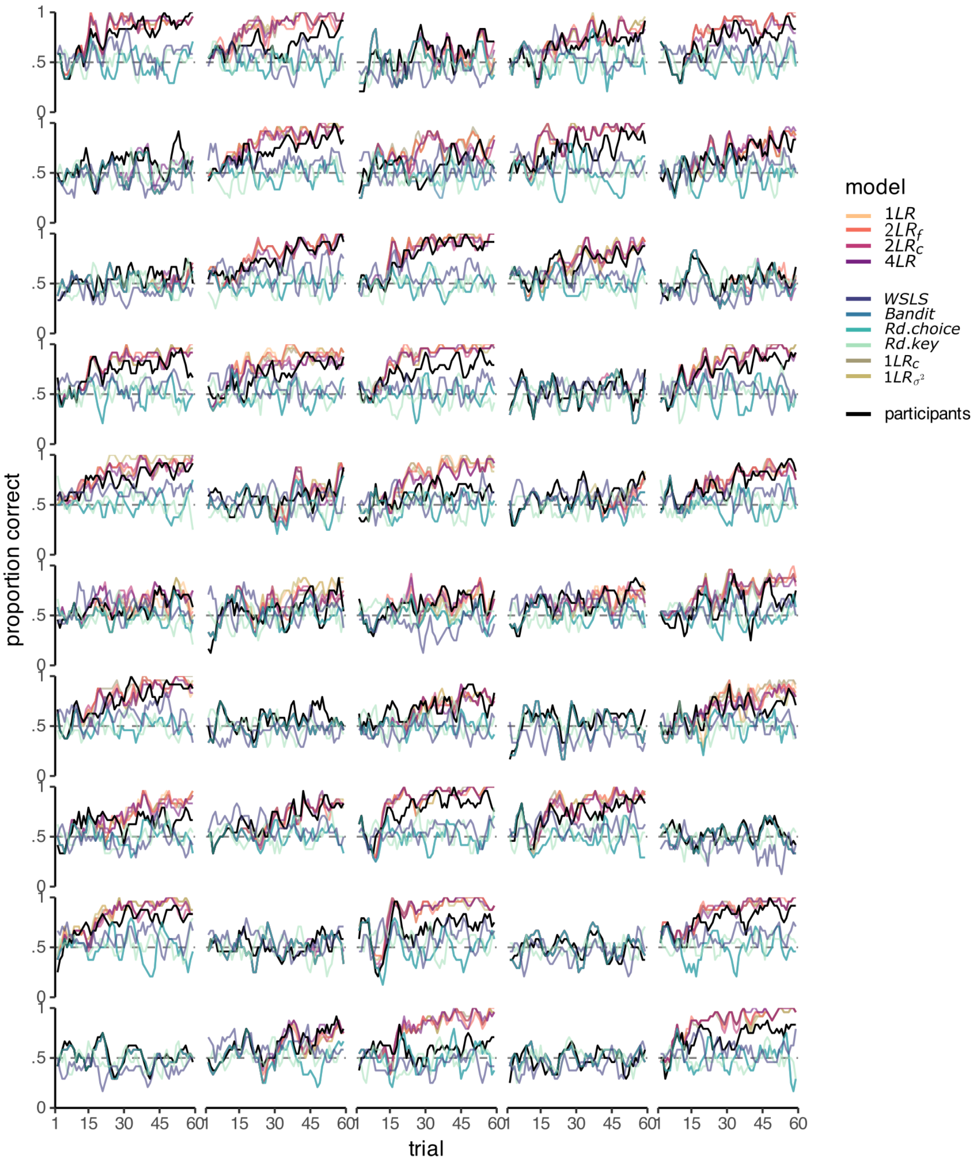


Figure B: **Individual participant and best fit model learning curves for experiment 2.** Participant learning curves overlaid with the model learning curves, simulated using the parameter values obtained from maximum likelihood fitting. Error bands are not shown for better visibility.

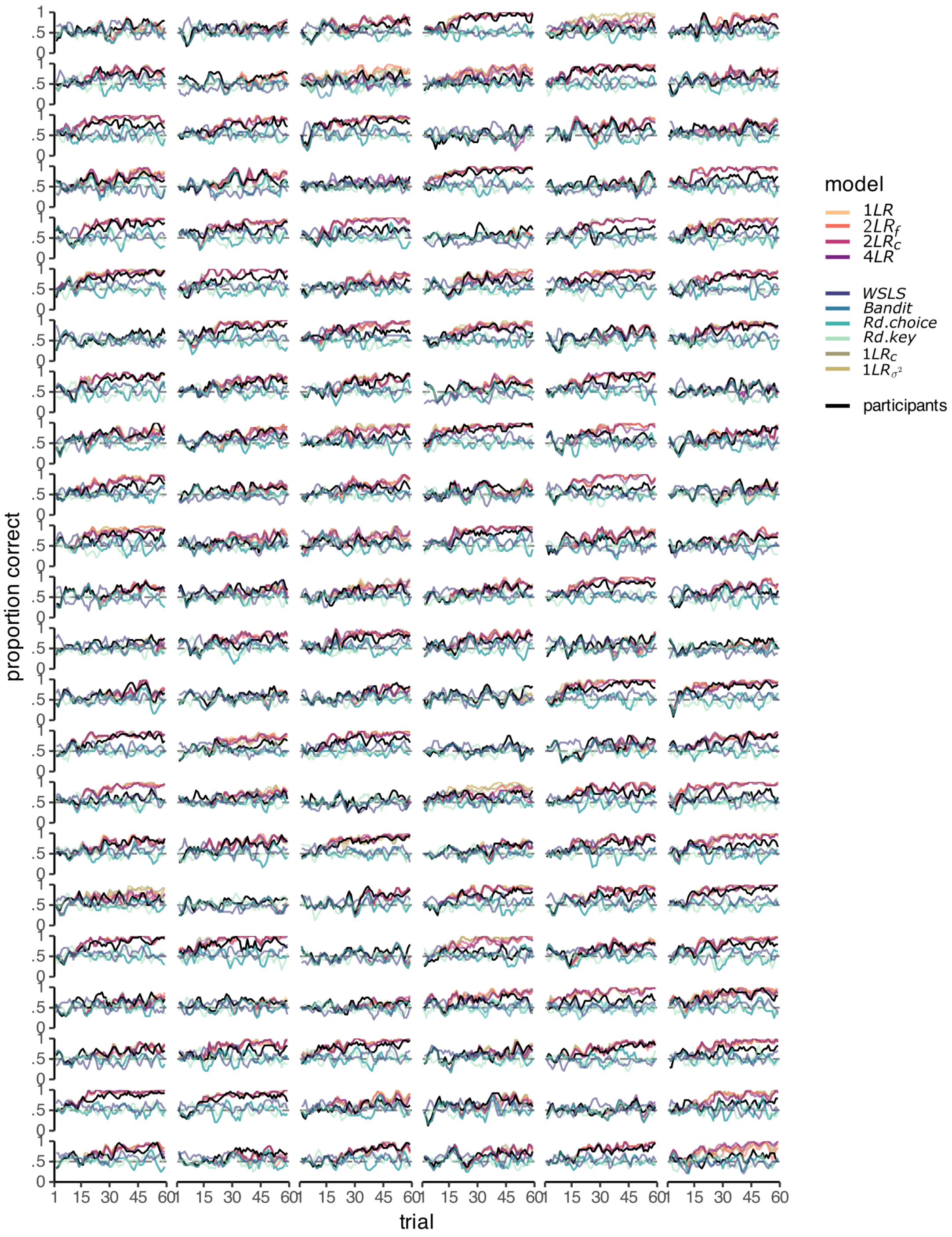


Figure C: **Individual participant and model learning curves for the replication.** Participant learning curves overlaid with the model learning curves, simulated using the parameter values obtained from maximum likelihood fitting. Error bands are not shown for better visibility.

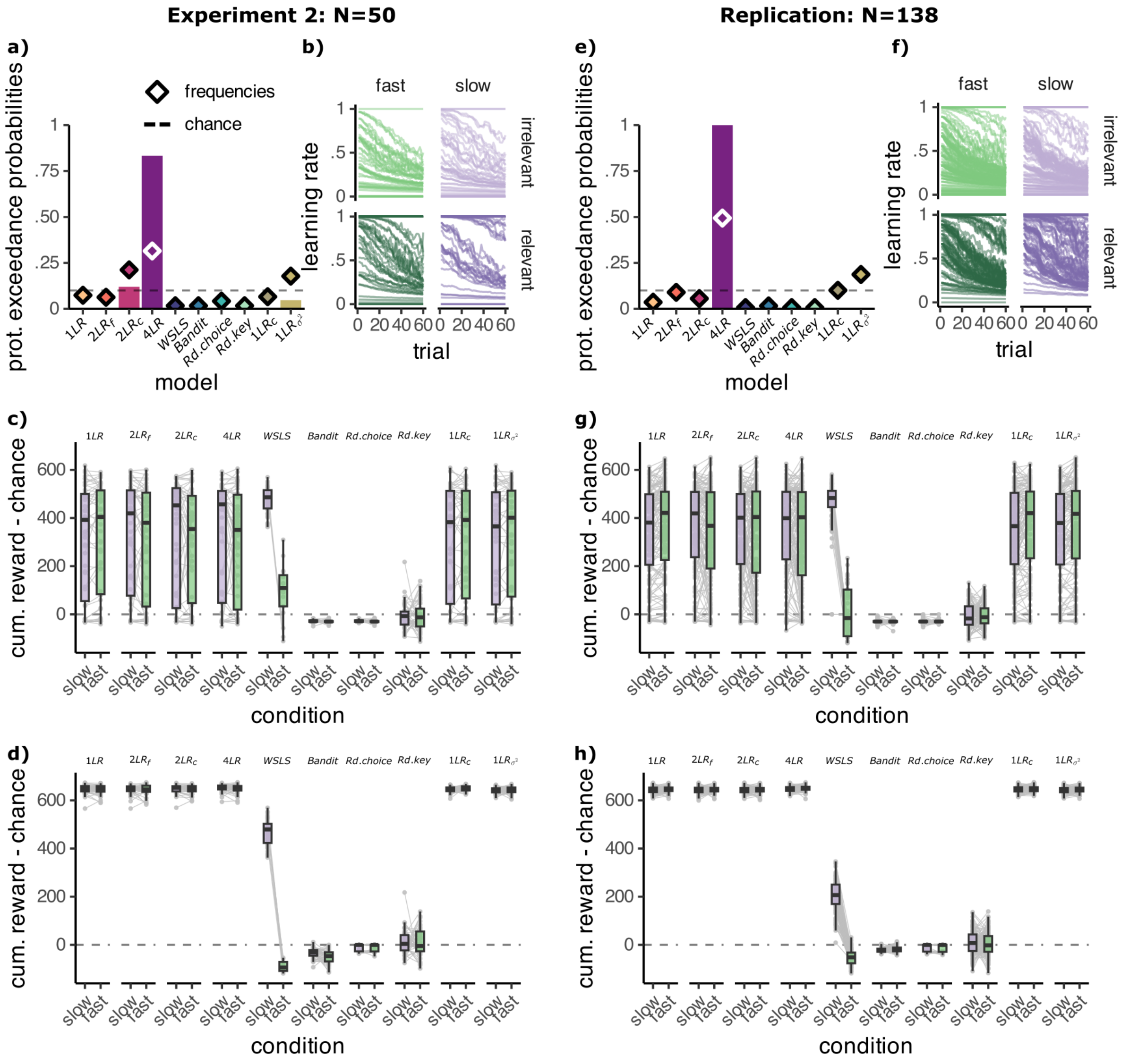


Figure D: **Model performance.** **A/E:** Exceedance probabilities plot including the models shown in the main paper and the two additional models, which allowed the exploration parameter $c$ ($1\text{LR}_c$) or the decision noise parameter $\sigma^2$ ($1\text{LR}_{\sigma^2}$) to vary by slow/fast condition. **B/F:** The development of the learning rates of the four learning rates model across the trials of the learning phase. The learning rates on the first trial were obtained through maximum likelihood fitting. With decreasing uncertainty about the stimulus values, the learning rates decreased. The learning rates could not increase. Each line is an individual participant, averaged across blocks. **C/D/G/H:** Cumulative reward relative to chance level obtained in the learning phase by the models in the same curriculum as participants using **C/G:** maximum likelihood parameters **D/H:** reward maximising parameters.

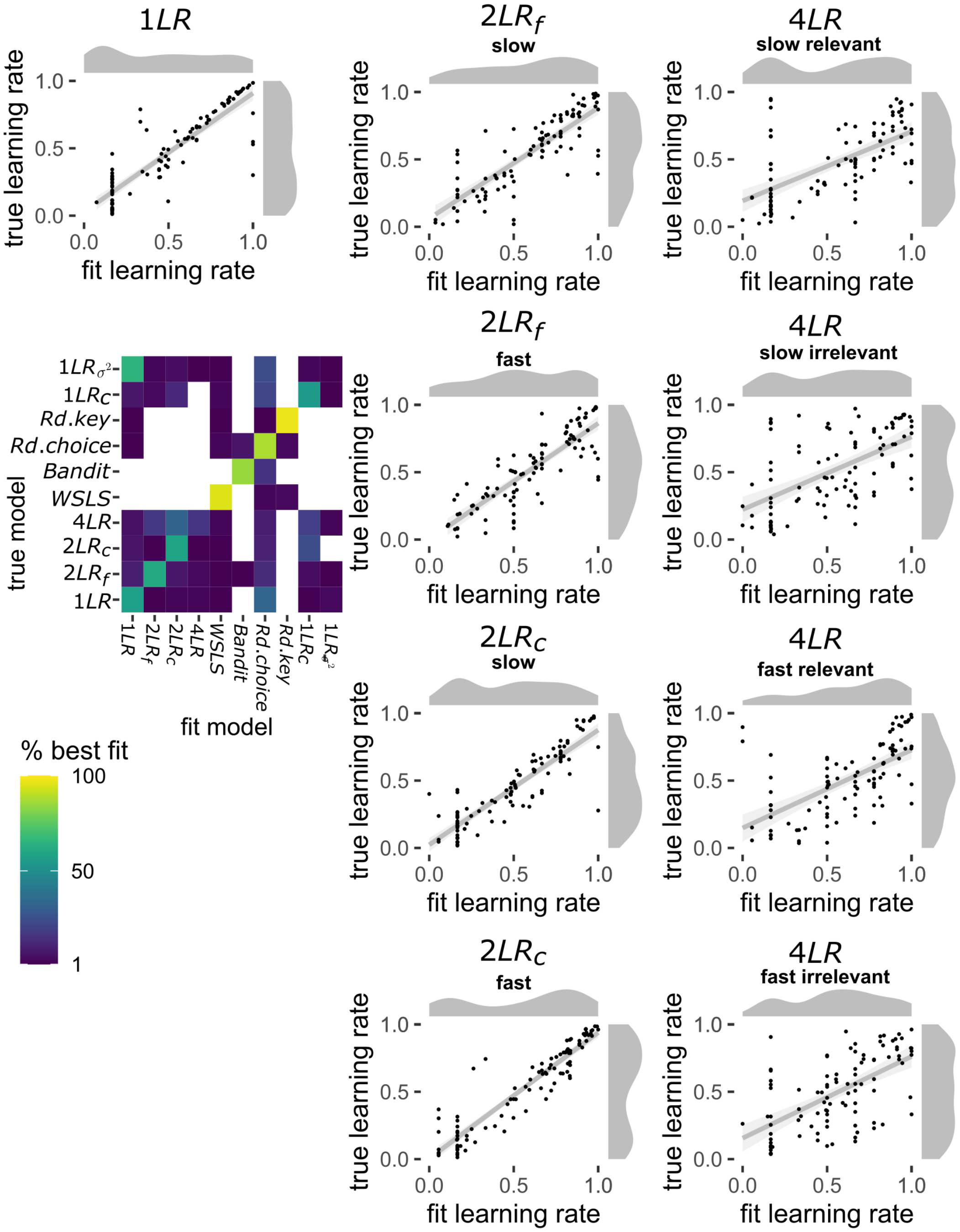


Figure E: **Parameter and model recovery.** Model fitting and comparison was performed as described in the Methods. 100 data-sets were simulated for each model using random parameter values. Dot plots show the relationship between the true learning rate and the learning rate obtained through maximum likelihood fitting. Distributions of both are shown in grey. Points are individual simulations. The matrix shows model recovery. True models are accurately identified in most simulations, except for the $1LR_{\sigma^2}$ model. The 4LR model is sometimes misidentified as the $2LR_c$ and to a lesser extent the $2LR_f$ model. This likely is because the 4LR model is a specialised case of these two models. If the four learning rates of the 4LR model are similar to each other, this model behaves similarly to the two-learning (2LR) rate models.

### List of Legends

**Fig A. Participant learning curves and meta analysis. A:** Participant learning curves for the slow and fast condition, showing the increase in the accuracy of choices across trials of the learning phase in the slow (purple) and fast(green) blocks. Each panel shows a different experiment sample. **B:** Results of the meta analysis of the cumulative reward in the learning phase, confirming consistently higher reward in the slow blocks. **C:** Results of the meta analysis of the accuracy in the test phase, showing no consistent advantage in generalisation in slow blocks. Plots produced with the `forest` function from the `meta` package in R [1].

**Fig B. Individual participant and best fit model learning curves for experiment 2.** Participant learning curves overlaid with the model learning curves, simulated using the parameter values obtained from maximum likelihood fitting. Error bands are not shown for better visibility.

**Fig C. Individual participant and model learning curves for the replication.** Participant learning curves overlaid with the model learning curves, simulated using the parameter values obtained from maximum likelihood fitting. Error bands are not shown for better visibility.

**Fig D Model performance. A/E:** Exceedance probabilities plot including the models shown in the main paper and the two additional models, which allowed the exploration parameter $c$ ($1LR_c$) or the decision noise parameter $\sigma^2$ ($1LR_{\sigma^2}$) to vary by slow/fast condition. **B/F:** The development of the learning rates of the four learning rates model across the trials of the learning phase. The learning rates on the first trial were obtained through maximum likelihood fitting. With decreasing uncertainty about the stimulus values, the learning rates decreased. The learning rates could not increase. Each line is an individual participant, averaged across blocks. **C/D/G/H:** Cumulative reward relative to chance level obtained in the learning phase by the models in the same curriculum as participants using **C/G:** maximum likelihood parameters **D/H:** reward maximising parameters.

**Fig E Parameter and model recovery.** Model fitting and comparison was performed as described in the Methods. 100 data-sets were simulated for each model using random parameter values. Dot plots show the relationship between the true learning rate and the learning rate obtained through maximum likelihood fitting. Distributions of both are shown in grey. Points are individual simulations. The matrix shows model recovery. True models are accurately identified in most simulations, except for the $1LR_{\sigma^2}$ model. The 4LR model is sometimes misidentified as the $2LR_c$ and to a lesser extent the $2LR_f$ model. This likely is because the 4LR model is a specialised case of these two models. If the four learning rates of the 4LR model are similar to each other, this model behaves similarly to the two-learning (2LR) rate models.
